# Elimination Of Aberrantly Specified Cell Clones Is Independent Of Interfacial Myosin II Accumulation

**DOI:** 10.1101/2021.05.14.444162

**Authors:** Olga Klipa, Fisun Hamaratoglu

## Abstract

Spatial organization of differently fated cells within an organ is essential and needs to be maintained during development. This is largely implemented via compartment boundaries that serve as barriers between distinct cell types. Biased accumulation of junctional non-muscle Myosin II along the interface between differently fated groups of cells contributes to boundary integrity and maintains its shape via increased tension [1–4]. Here we test whether interfacial Myosin driven tension is responsible for the elimination of aberrantly specified cells that would otherwise compromise compartment organization. To this end, we genetically reduce Myosin II levels in three different patterns: in both wild-type and misspecified cells, only in misspecified cells and specifically at the interface between wild-type and aberrantly specified cells. We find that recognition and elimination of aberrantly specified cells do not rely on tensile forces driven by interfacial Myosin cables. Moreover, apical constriction of misspecified cells and their separation from wild type neighbors occurs even when Myosin level is greatly reduced. Thus, we conclude that the forces that drive elimination of aberrantly specified cells are largely independent of Myosin II.

## INTRODUCTION

Spatial organization of differently fated cells is central to the development of multicellular organisms. Correctly organized cells are vital in ensuring the formation of fully functional body structures and organs. Thus, cells with inappropriate positional identity, which occasionally arise due to sporadic mutations or chromatin defects, are detrimental for the whole organism. Fortunately, such cells can be detected and effectively removed from the tissue [4–7]. However, the mechanism by which this happens is largely unknown. An ideal model to address this question is the *Drosophila* wing imaginal disc, because of its relatively simple and well described patterning events and its readily available mosaic techniques [8].

In *Drosophila*, wing disc cells are organized into four compartments (anterior (A), posterior (P), dorsal (D) and ventral (V)) separated by lineage boundaries. The cellular identity of each compartment is defined by the restricted expression of selector genes. Expression of *engrailed* and *invected* grants cells a posterior identity, whereas their absence marks cells for an anterior fate. Likewise, *apterous (ap)* expression defines dorsal cells whereas its absence specifies ventral cells. The interactions between anterior and posterior cells as well as that between dorsal and ventral cells firstly lead to local activation of short-range signaling molecules, and secondly, secretion of long-range morphogens Decapentaplegic (Dpp) and Wingless (Wg) along anteroposterior (AP) and dorsoventral (DV) boundaries, respectively. Therefore, the compartment boundaries act both as fences between different cell populations, preventing their mixing, and as organizing centers for further patterning events [9, 10].

Conceptually, two mechanical properties keep cell populations apart: differences in cohesive strength between interacting cell types (differential adhesion), and local increase of junction tension along the interface (interfacial tension) [10–16]. The ways in which differential adhesion and interfacial tension are genetically encoded and controlled by upstream signaling events have been the subject of intense research. In the case of the D/V boundary, both processes contribute to boundary formation and maintenance. Ap target genes *capricious, tartan, fringe, serrate, delta* and *bantam*-microRNA were shown to play key roles in the process [17–21]. Transmembrane proteins Capricious and Tartan are dorsally expressed at the time of D/V boundary formation and contribute to cell segregation [17]. They are thought to function as ligands to a yet to be discovered dorsal specific receptor [22]. Thus, the initial separation of dorsal and ventral cells is achieved via early dorsal expression of Capricious and Tartan [18]. Once the boundary is formed, its maintenance depends on Notch (N) activity [18, 19, 21, 23–26]. Ap induces the expressions of the N ligand Serrate and its modulator Fringe in dorsal cells and restricts Delta expression to the ventral cells ensuring a stripe of N signaling along the D/V boundary [19, 20, 27] which is thought to increase the cell bond tension [26]. It was shown that actomyosin filaments are enriched along the DV boundary [1, 25] and, cortical tension is higher at the boundary than elsewhere in the compartments [3, 28]. Actomyosin enrichment and elevated tension at lineage boundaries ensure that differently fated cells are kept separate and straight boundaries are maintained [2, 3, 28–30].

Crucially, if clones of cells mutant for a selector gene arise in a compartment where this gene is expressed, the clones are either relocated to the opposite compartment or eliminated from the tissue completely [7, 31–34]. Cell sorting (relocation) and elimination were also reported for clones that ectopically express a range of other fate specifying transcription factors (including cubitus *interruptus, vestigial, homothorax, Iroquois Complex, spalt* and *optomotor-blind*) [4, 35–38]. Similarly, cell clones with abnormal levels of Wg or Dpp signaling were shown to form cysts or undergo apoptosis [6, 39–44]. Altogether these numerous observations demonstrate the ability of a tissue to identify and remove, or sort out misspecified cells. However, the mechanism by which this is achieved remains unclear.

In this study, we focus on cells with aberrant dorsoventral identity. Cells that lack Apterous activity are eliminated from the dorsal compartment of the wing disc, whereas cells that ectopically express Apterous are removed from the ventral part (Fig. 1A) [7, 34]. There are at least three mechanisms involved in the clearance: relocation to the identity-appropriate compartment, apoptosis, and basal extrusion [7]. Interaction of *ap*-expressing and *ap*-non-expressing cells at the boundary of misspecified clones induces ectopic activation of Notch and Wingless signaling. The clone boundaries become smooth, suggesting its separation from the surrounding wild-type cells [18]. This greatly resembles the process at the DV boundary. Altogether these observations raise the question of whether the events involved in the formation of compartment boundaries during normal development also play a central role in cell elimination. Accordingly, modulation of Notch signaling or reestablishment of adhesive properties allows aberrantly specified cells to remain in the tissue [18, 34]. Moreover, it was reported that Myosin II accumulation and increased tension at the DV boundary depends on Notch activity [1, 26]. Thus, separation of cells with incorrect dorsoventral identity and their subsequent elimination could be mediated by increased tension, driven by actomyosin contractility along clone borders. Importantly, this was proposed to be a general mechanism by which the tissue identifies and responds to the presence of misspecified cells [4]. Indeed, Bielmeier and colleagues reported accumulation of actomyosin filaments at the interface between differently fated cells upon misexpression of a range of fate specifying genes [4]. The beauty of this model is in that it can account for the elimination of any misspecified clone from the tissue regardless of the clone’s genetic identity, and the particular events that culminates in actomyosin accumulation.

**Figure 1:**
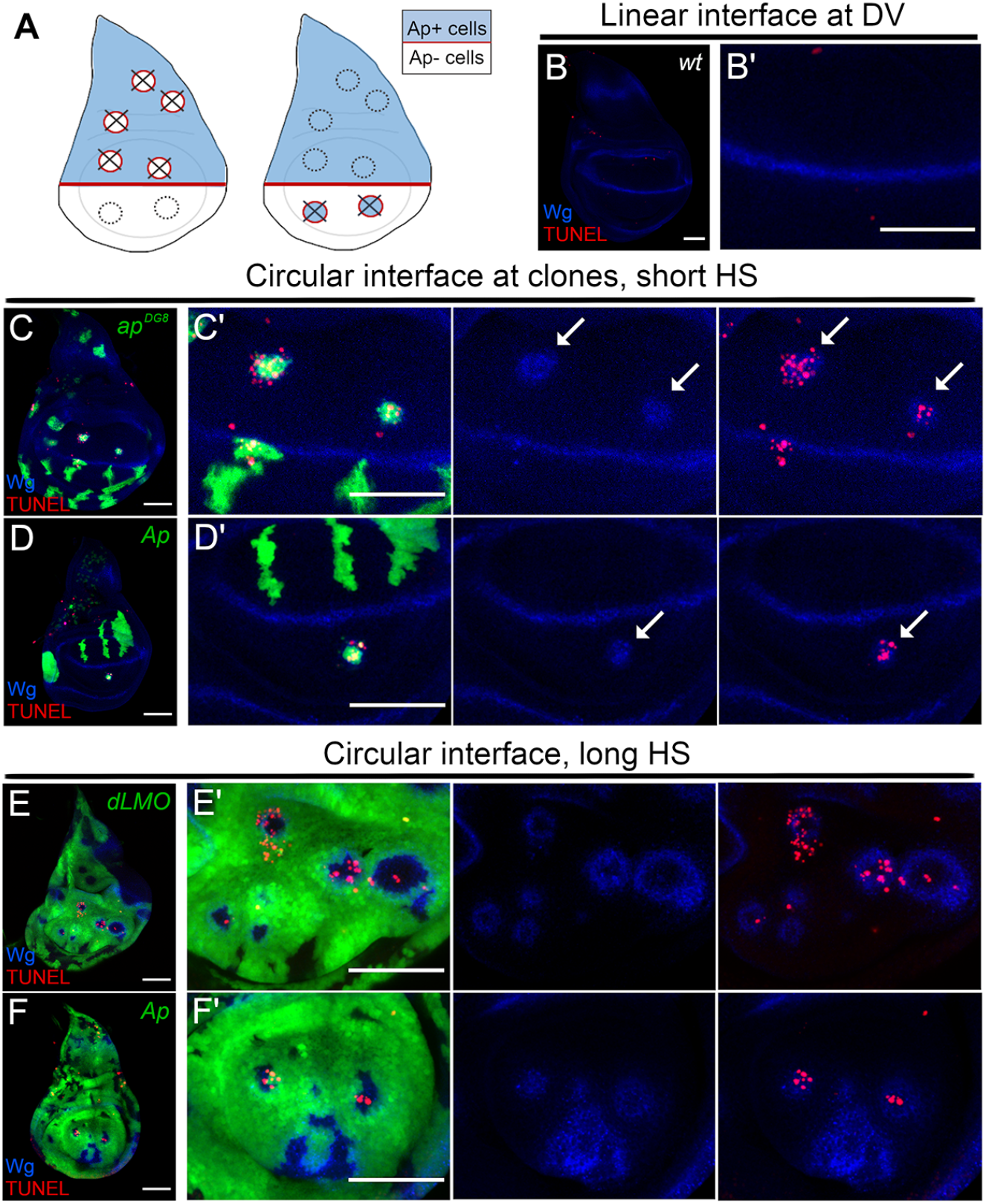
The majority determines the correct cell fate in the wing disc and the cells of “wrong” identity undergo apoptosis. **A)** Schematics of third instar wing discs. Ap (blue) is expressed in the dorsal compartment and Wg (red) expression marks the D/V boundary. Removal of Ap function from dorsal compartment (left) or ectopic Ap expression in the ventral compartment (right) leads to Wg expression at the interface of Ap+ and Ap-cells and eventual elimination (crosses). **B-B’)** Third instar wing disc (B) and its pouch region (B’) showing Wg antibody staining in blue and TUNEL in red. Wg expression at the boundary reveals a straight interface between Ap-expressing dorsal cells and ventral cells without Ap. **C-D’)** Wing discs bearing GFP-expressing (green) clones that are mutant for *ap^DG8^* (C) or ectopically expressing Ap (D) and their pouch regions (C’, D’). All panels show Wg antibody staining in blue, and TUNEL in red. Arrows point to circular ectopic boundaries associated with misspecified clones that are undergoing elimination. **E-F’)** A long heat-shock generates discs that are almost entirely composed of GFP-positive (green) *dLMO* (E) or Ap-expressing (F) cells. (E’, F’) Pouch regions of the discs shown in (E, F) with separate channels showing Wg (blue) and TUNEL (red) stainings. Wild-type islets trapped in between overexpressing cells undergo apoptosis. All scale bars represent 50μm. Dorsal is up, anterior is to the left in all panels.

Here, we examine the role of non-muscle Myosin II, the main regulator of contractile forces in a cell, in elimination of cell clones with inappropriate dorsoventral identity. To address this question, we interfered with Myosin II activity in three different patterns: in the whole disc, inside the clone and at the clone boundary. We then examined the clones’ topology and elimination efficiency. Surprisingly, we found that both clone elimination and its separation from surrounding cells do not rely on Myosin II.

## RESULTS

### Apposition of differently fated cells induces apoptosis when the contact interface is circular but not when it is linear, ensuring the removal of the underrepresented cell population

Both dorsal and ventral cell populations are healthy and viable on their own. The interaction of Apterous (Ap)-expressing cells (dorsal identity) and Ap-non-expressing cells (ventral identity) causes activation of boundary signaling (Notch and Wingless). However, the outcome of this event at the DV compartment boundary is different to that around mispositioned clones. The activation of N / Wg signaling along the compartment boundary does not lead to cell death (Fig. 1B-B’). It is a physiological condition that is necessary for proper patterning. In contrast, the interaction of misspecified cell clones with their wild-type (wt) neighbors induces apoptosis as revealed by the TUNEL assay. Both *ap* mutant cells in the dorsal compartment and Ap-expressing cells in the ventral compartment show strong induction of apoptosis (Fig. 1C-D’, arrows). Thus, the apoptosis observed purely relies on cell-cell interactions, and does not depend on cell identity or the clone’s topological localization in the disc. To further test this idea, cell clones expressing either *dLMO* (negative regulator of Ap activity) or *Ap* were induced using a long heat-shock scheme to allow these cells to occupy almost the whole wing disc. The remaining wt cells, GFP negative, were highly underrepresented (Fig. 1E-F’). The wing discs with such clones did not have regular DV boundaries, as revealed by Wg staining. Instead, they contained boundary signal in circles that were formed around the wt islands (Fig. 1E-F’). Importantly, the interaction of differently fated cells in this scenario also resulted in cell death. However, this time apoptosis was preferably associated with the wt cell population (Fig. 1E-F’). This suggests that among two differently fated cell populations that interact with each other, the underrepresented one is designated for elimination regardless of its identity. When one cell population is underrepresented, the boundary it forms with surrounding neighbors is circular. In contrast, the compartment boundary, which is formed between relatively big and similarly sized cell populations, is linear. The former is associated with apoptosis and elimination whereas no cell death is induced in the latter. Thus, it is possible that the presence of relatively small groups of cells that disrupt global pattern (misspecified cells) could be detected by a circular interface. The underlying mechanism as to how interface shape determines the output could be attributed to interface contractility. According to this model, increased contractility around relatively small cell clusters causes irresistible mechanical stress (compression and apical constriction) in this encircled groups of cells, eventually leading to induction of apoptosis, and subsequent elimination [4].

### Evidence for interface contractility between Ap-positive and Ap-negative cells

To investigate whether the interface between differently fated cell populations in our scenario has elevated tension, we analyzed circularity of the interfaces at wt GFP-marked clones surrounded by wt cells (Fig. 2A), *ap* mutant clones located in the dorsal compartment (Fig. 2B) and Ap-positive wt islands surrounded by *dLMO*-expressing cells (Fig. 2C). The interface between differently fated cells was more circular compared to the interface between cells of the same identity (Fig. 2D). To test whether the enclosed cells outlined by highly circular boundaries experience compression, we measured the apical areas of individual cells from both sides of the interface, within the clones and outside. The cell membranes were revealed by DE-Cadherin staining and the analysis was done using Epitools [45]. The cells of wt clones had similar apical areas as the surrounding wt cells (Fig. 2A, heatmap and 2E). In contrast, the apical areas of the *ap^DG8^* cells and those of cells in wt islands were much smaller compared to that of the surrounding cells (Fig. 2B-C, heatmaps and 2E). Importantly, the surrounding cells had comparable apical areas in all three scenarios (Fig. 2E, blue bars). Such apical constriction of underrepresented cells could be easily explained by interface contractility. Traditionally, contractility and high tension are associated with enrichment of actomyosin filaments. Therefore, we analyzed F-actin (using Phalloidin staining) and non-muscle Myosin II (using a *sqh-GFP* reporter) distribution at misspecified cell clones, and wt islets surrounded by cells of different identity. Both F-actin and Myosin II accumulated at adherens junctions along the borders of many (but not all) *dLMO* clones that remained in the dorsal compartment (Fig. 3A-B’). Interestingly, in some cases elevated levels of Myosin were observed not only along the interface junctions but also at the junctions inside the misspecified clones (Fig. 7A-A’). The actomyosin enrichment was even more prominent along the interfaces between wt islands and *dLMO*-expressing cells generated by long heat-shock. Very thick F-actin and Myosin cables were observed in nearly all cases (Fig. 3C-D’). Altogether these observations provide evidence of increased contractility along the interface and are in favor of the mechanical stress model. According to this model, the elimination of misspecified cells is due to their deformation mediated by actomyosin accumulation and increased contractility at the interface. If this is the case, then depletion of Myosin would be enough to abolish the elimination. In order to test this hypothesis, we designed experiments where the level of Myosin was reduced in the whole disc, within the misspecified clones and specifically at the clone boundary.

**Figure 2:**
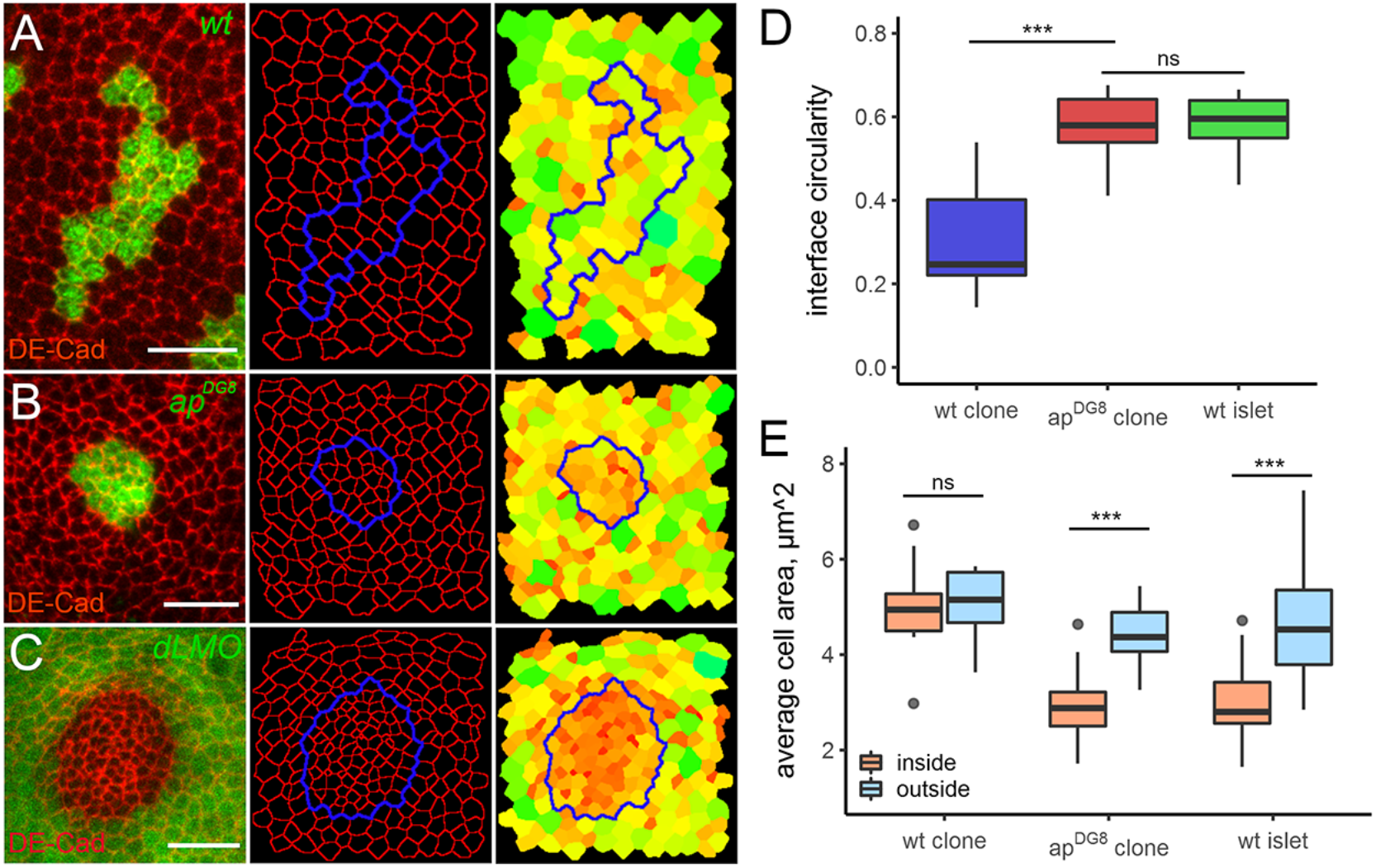
Cells undergo apical constriction within the patches of minority identity. **A-C)** Representative examples of GFP-marked (green) wild-type (A) or *ap^DG8^* mutant (B) clones, and a GFP-negative wild-type islet (C) in a *dLMO*-expressing disc. DE-Cad (red) staining was used to reveal the apical cell outlines in Epitools. Clone boundaries are shown in blue. The right panels show heat-maps of apical areas, from smallest (0.2μm^2^, dark red) to largest (12μm^2^, light green). **D-E)** Quantifications of interface circularity (D) and average cell area (E) inside (orange) and outside (blue) from 11 wild-type, 20 *ap^DG8^* clones and 25 wild-type islets trapped in *dLMO* expressing discs. All scale bars represent 10μm.

**Figure 3:**
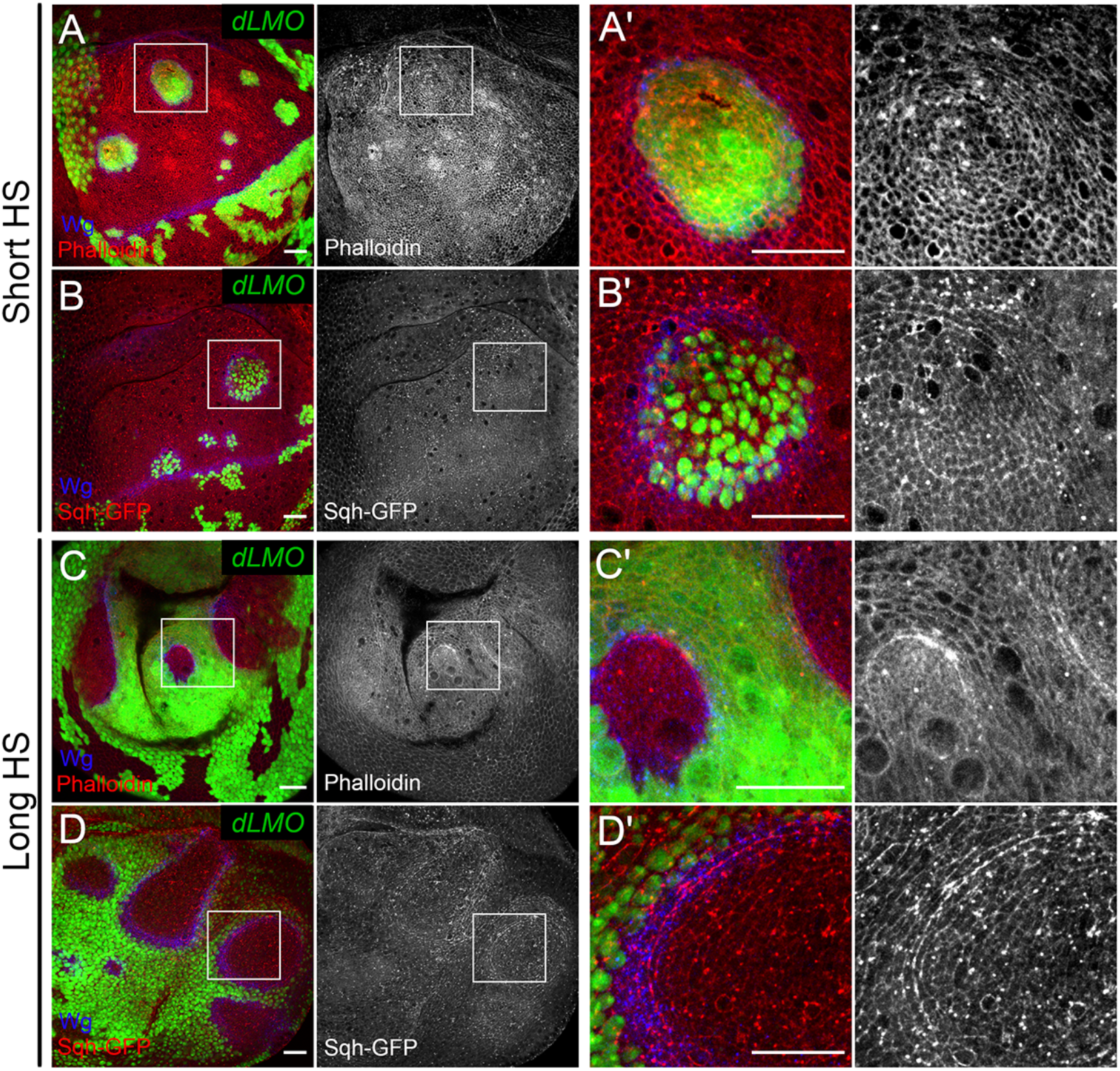
Actomyosin network is enriched at the interface between cell populations of different identities. **A-B)** Wing discs with *dLMO*-expressing cells (green), stained for Wg (blue) and Phalloidin (red, grey) (A-A’) or carrying the Sqh-GFP transgene (red, grey) (B-B’). **C-D)** Wing discs subjected to long heat-shock generating wild-type islets (non-green) surrounded by *dLMO*-expressing cells (green). Wg is shown in blue; red in overlay or single grey channels show Phalloidin (C-C’) or Sqh-GFP (D-D’). A’-D’ show the zoomed in versions of the white boxes in A-D, respectively. All scale bars represent 20μm. Dorsal is up, anterior is to the left in all panels.

### Misspecified clones induced in Myosin mutant wing discs are still eliminated

Non-muscle Myosin II is a vital molecule and its function is highly important for development. That is why animals bearing a strong Myosin mutation are not viable. However, some combinations of hypomorphic alleles of Myosin heavy chain *zipper (zip)* are less harmful and flies can survive until later stages. It was shown that *zip^Ebr^/zip^2^* is the strongest combination that allows recovery of third instar larvae [1]. We assessed Myosin levels in the *zip^Ebr^/zip^2^* wing discs using antibodies against phosphorylated form of Myosin light chain (p-MLC), which is an indicator of assembled active form of the Myosin complex. In *zip^Ebr^/zip^2^* wing discs, Myosin levels were greatly reduced when compared to wt discs, yet some residual junctional Myosin was observed (Fig. 4A, 4B). Interestingly, the mutant discs were similar in size to wt discs of the same developmental stage (Fig. 4C, 4E). To evaluate the elimination efficiency of misspecified clones in Myosin mutant background we generated GFP-labeled wt and *dLMO*-expressing clones in both wt and *zip^Ebr^/zip^2^* imaginal discs at 58h after egg laying (AEL) and examined clone recovery in the dorsal compartment 60h later. As expected, *dLMO* clones were highly underrepresented in the dorsal part of the disc compared to wt clones (Fig. 4C-D and 4G). The area of dorsal compartment occupied by wt clones in wt and *zip^Ebr^/zip^2^* backgrounds were comparable (Fig. 4C, 4E and 4G). The recovery rate of *dLMO* clones generated in *zip* mutant discs was not significantly different from *dLMO* clones generated in wt discs. In both cases misspecified clones were eliminated efficiently (Fig. 4D, 4F and 4G). Next, we examined whether the *zip^Ebr^/zip^2^* background affects the increased interface circularity and apical constriction associated with the aberrantly specified clones. Surprisingly, we found that dorsal *dLMO* clones in Myosin mutant discs had very smooth borders (Fig. 4K) and their circularity did not differ from that of dorsal *dLMO* clones generated in a wt disc (Fig. 4I, 4K and 4L). Moreover, like *dLMO*-expressing cells induced in wt discs (Fig. 4I), *dLMO*-expressing cells in *zip* mutant discs (Fig. 4K) had smaller apical areas compared to the surrounding cells (Fig. 4M). As expected, the apical cell areas of GFP labeled clones in either wt (Fig. 4H) or Myosin mutant backgrounds (Fig. 4J) were not distinct from their surrounding neighbors (Fig. 4M). Thus, we conclude that reduction of Myosin using a *zip^Ebr^/zip^2^* allelic combination does not result in a relaxation of tension along the borders of misspecified clones nor does it prevent their elimination.

**Figure 4:**
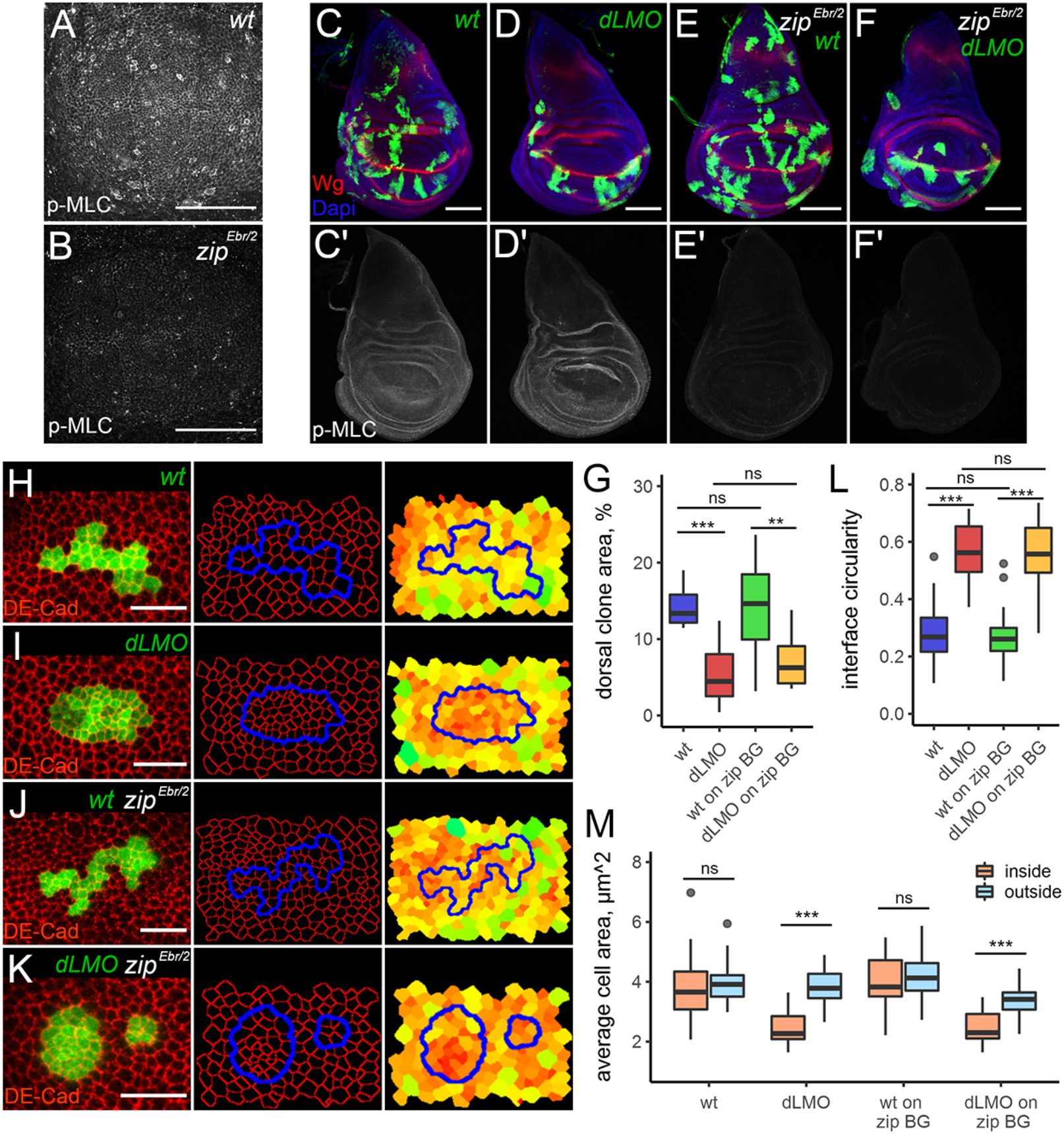
A reduction in Myosin activity has no effect on interface circularity, apical constriction, and *dLMO*-expressing cell clone elimination. **A-B)** Phospho-Myosin Light Chain (p-MLC) staining (grey) is reduced in *zip^Ebr/2^* mutant background (B), compared to wild-type (A). The samples were processed in parallel. The scale bars in A and B are 50μm. **C-F’)** Wing discs containing control (C and E) or *dLMO*-expressing clones (D and F) in a wild-type (C, D) or *zip^Ebr/2^* mutant background (E, F). C’-F’) show p-MLC staining (grey) alone of the corresponding panels above. Scale bars are 100μm in (C-F). Dorsal is up, anterior is to the left in all panels. **(H-K)** Representative examples of GFP-marked (green) wild-type (H, J) or *dLMO*-expressing (I, K) clones in a wild-type (H, I) or *zip^Ebr/2^* mutant background (J, K). DE-Cad (red) staining was used to reveal the apical cell outlines in Epitools. Clone boundaries are shown in blue. The right panels show heatmaps of apical areas, from small (red) to large (green). Scale bars in (H-K) are 10μm. **G)** Quantification of dorsal clone area in indicated genotypes. Minimum 12 discs were analysed per genotype. **L-M)** Quantifications of interface circularity (L) and average cell area (M) inside (orange) and outside (blue) from minimum 30 clones for each genotype.

### Myosin II depletion inside the misspecified clones has a mild effect on clone recovery

Next, we obstructed Myosin II function within misspecified clones by knocking down the regulatory light chain of Myosin II, *spaghetti squash* (*sqh*). The *UAS-sqh-RNAi (sqhRi)* line used is very effective in depleting Myosin as revealed by phospho-Myosin staining. The Myosin levels in cell clones expressing *sqhRi* dramatically decreased, with no residual Myosin cables detected at the junctions of these cells (Fig. 5A-A’). Interestingly, many *sqhRi* clones were dispersed in appearance, where some clonal cells disengaged from their siblings and became fully surrounded by wt cells (Fig. 5A, arrowhead and Fig. S1A-A’). To analyse this effect quantitatively, we introduced two parameters: *the mixing index*, defined as the number of wt cells in direct contact with one clonal cell at the frontline (Fig. S1B-B’), and *the contact length*, the portion of clone perimeter per clonal cell at the frontline (Fig. S1B’’). Both the mixing index and the contact length of *sqhRi* clones were significantly higher than that of wt clones (Fig. S1C-D). Thus, *sqhRi* impairs clone integrity and allows clonal cells to split apart by cell divisions and rearrangements more easily. These observations suggest that knocking down Myosin II by *sqhRi* compromises tensile forces at cell junctions. To assess the effect of *sqhRi* on elimination of misspecified clones, wt, *dLMO*, *sqhRi* and *dLMO + sqhRi* clones were induced at 58h AEL and clone recovery in the dorsal disc was measured 60h later. The *dLMO*-expressing clones were eliminated as expected (Fig. 5B-C, 5F) and the recovery of *sqhRi* cells was comparable to that of wt cells (Fig. 5D and 5F). Interestingly, co-expression of *dLMO* with *sqhRi* also resulted in efficient clone elimination from the dorsal compartment (Fig. 5E and 5F). However, the recovery rate of such clones was slightly higher when compared to the one of clones expressing *dLMO* alone (Fig. 5F). Next, we examined how *sqhRi* co-expression affects tension along misspecified clone boundaries. As mentioned above, *sqhRi*-expressing clones easily split apart and have a high mixing index and long contact lengths (Fig. S1). However, this increased mixing tendency of *sqhRi* cells was not reflected in our measurements of interface circularity; *sqhRi* clones did not differ from wt clones in terms of circularity (Fig.5G, 5I and 5K). This is likely due to the fact that we only analyzed clones consisting of ≥10 cells for their circularity. If all the dispersed cells could be included in our analysis as part of the bigger clone they separated from, lower circularity would be yielded for *sqhRi*-expressing clones. As expected, *dLMO* clones round up efficiently and have very smooth borders (Fig. 5H and 5K). Clones co-expressing *dLMO* and *sqhRi* were slightly less circular compared to those expressing *dLMO* alone (Fig. 5H, 5J and 5K). Moreover, in some instances, *dLMO + sqhRi* clonal cells appeared to be detached from the main clone (Fig. 5J, arrowhead). Nevertheless *dLMO + sqhRi* clone borders were much smoother than that of either wt or *sqhRi* clones (Fig 5G, 5I, 5J and 5K). In addition, *sqhRi* co-expression did not change the mixing index or the contact length of misspecified clones (Fig. S1C, S1D). Notably, the *sqhRi* clones were highly heterogeneous in cell size, hosting both very small and very big cells (Fig. 5I, heatmap). However, the average apical areas of *sqhRi*-expressing cells were similar to those of the surrounding wt cells (Fig. 5I, heatmap and 5L). Strikingly, the apical areas of cells within *dLMO* as well as *dLMO + sqhRi* clones, were much smaller than that of surrounding cells (Fig. 5H, 5J, heatmaps and 5L). Thus, co-expression of *sqhRi* did not prevent the apical constriction of misspecified cells. We conclude that although depletion of Myosin II in misspecified clones leads to interface relaxation and slightly increases clone recovery, the effect is too mild to grant Myosin a central role in elimination of misspecified cells.

**Figure 5:**
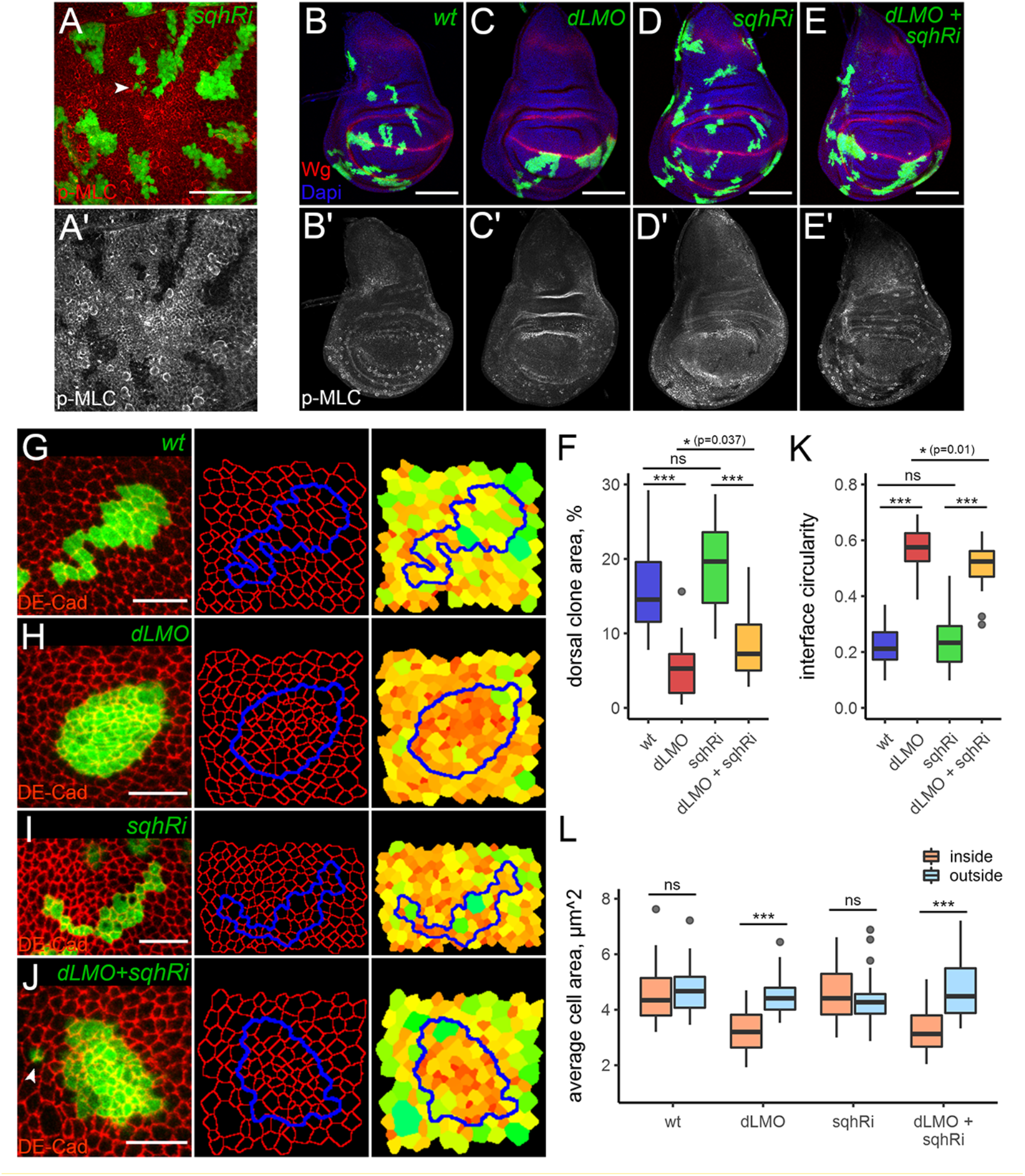
Knocking down Myosin light chain within *dLMO*-expressing clones mildly influences clone recovery and circularity. **A-A’)** Phospho-Myosin Light Chain (p-MLC) staining (red in A and grey in A’) is greatly reduced in *sqh-RNAi*-expressing cell clones (green). Arrowhead in (A) points to dispersed cells. The scale bar in (A) is 50μm. **B-E’)** Wing discs containing control (B), *dLMO*-expressing (C), *sqh-RNAi*-expressing (D), or *dLMO* and *sqh-RNAi*-expressing clones (E). B’-E’) show p-MLC staining (grey) alone of the corresponding panels above. Scale bars are 100μm in (B-E). Dorsal is up, anterior is to the left in all panels. **(G-J)** Representative examples of GFP-marked (green) wild-type (G), *dLMO*-expressing (H), *sqh-RNAi*-expressing (I), or *dLMO* and *sqh-RNAi*-expressing clones (J) clones. DE-Cad (red) staining was used to reveal the apical cell outlines in Epitools. Clone boundaries are shown in blue. The right panels show heatmaps of apical areas, from small (red) to large (green). Scale bars in (G-J) are 10μm. **F)** Quantification of dorsal clone area in indicated genotypes. Minimum 15 discs were analysed per genotype. **K-L)** Quantifications of interface circularity (L) and average cell area (M) inside (orange) and outside (blue) from minimum 30 clones for each genotype.

### No evidence that Myosin II depletion along boundaries of misspecified clones prevents their elimination

The approach described above targeted Myosin in clonal cells only, without affecting its expression in flanking wt cells. Arguably both sides can contribute to Myosin enrichment and tension observed along clone boundaries. We therefore sought to design an experiment, where we reduce Myosin levels from both sides of the clone boundary. We took advantage of the fact that interaction of Ap-positive and Ap-negative cells results in Wg expression in flanking cells from both sides of the boundary. Thus, a *wg::Gal4* construct [46] was utilized to drive expression of *sqhRi* at the boundaries of misspecified cells. First, we analyzed how the expression of *sqhRi* in the Wg domain affects DV boundary of an otherwise wt wing disc (*wg>sqhRi*). The junctional Myosin levels were highly reduced at the DV boundary of such discs in comparison to that of control discs (w*g>GFP*), as revealed by p-MLC staining at the apical side (Fig. S2A-A’ and S2D-D’). The DV boundaries of *wg>sqhRi* discs appeared to be narrower than that of *wg>GFP* discs, and were occasionally disrupted, as revealed by the GFP signal (Fig. S2B-C’ and S2E-F’). Crucially, *wg>sqhRi* discs were smaller than the controls, suggesting some developmental delay (Fig 6A, B and 6E, see below). We induced *ap^DG8^* cells and analysed their recovery rates in both *wg>sqhRi* and *wg>GFP* discs. The clones were generated at 58h AEL and analyzed 60h later (day 5 AEL). The presence of *ap* mutant clones in the dorsal discs was revealed by the ectopic GFP expression induced around the clones (Fig. 6A-C). The number of remaining misspecified clones in the dorsal pouch of *wg>sqhRi* discs was slightly, but not significantly, higher than that in the dorsal pouch of *wg>GFP* discs. On average 2 misspecified clones per dorsal pouch remained in *wg>sqhRi* discs (Fig. 6A-B and 6D). However, *wg>sqhRi* larvae experienced some developmental delay, resulting in smaller discs at the specified time-point (Fig. 6E). To confirm that *ap* mutant clones undergo elimination in those discs, we also dissected larvae on day 6 AEL (84h after heat-shock). We found that on day 6 almost no misspecified clones remained in the dorsal pouch of *wg>sqhRi* discs (Fig. 6D). This suggests that despite the reduction of Myosin II along the clone boundary, the elimination of misspecified clones still takes place. Next, using Ap staining to define clone outlines, we analyzed the circularity of misspecified clones. Note that Ap, being a transcription factor, predominantly localizes to the nucleus. Therefore, we were unable to analyze clone circularity at the level of the adherens junctions (as in previous cases); instead, we measured it more basally. To ensure that more misspecified clones are retained for analysis, we generated the clones slightly later, at 68h AEL, and examined them 50h later. We noted that *wg::Gal4* drives expression of GFP in flanking cells as far as 2-3 cells on either side of the interface (Fig. 6F, 6G). In most cases, due to the small size of the misspecified clones, all clonal cells appeared to be GFP-positive. Surprisingly, the clones in the control (*wg>GFP*) and the experimental (*wg>sqhRi*) discs were both round, with very similar circularity (Fig. 6F’, 6G’ and 6H). Altogether, this data does not support the model in which the central role of clearance of aberrantly specified clones is attributed to Myosin-driven contractility at the clone interface.

**Figure 6:**
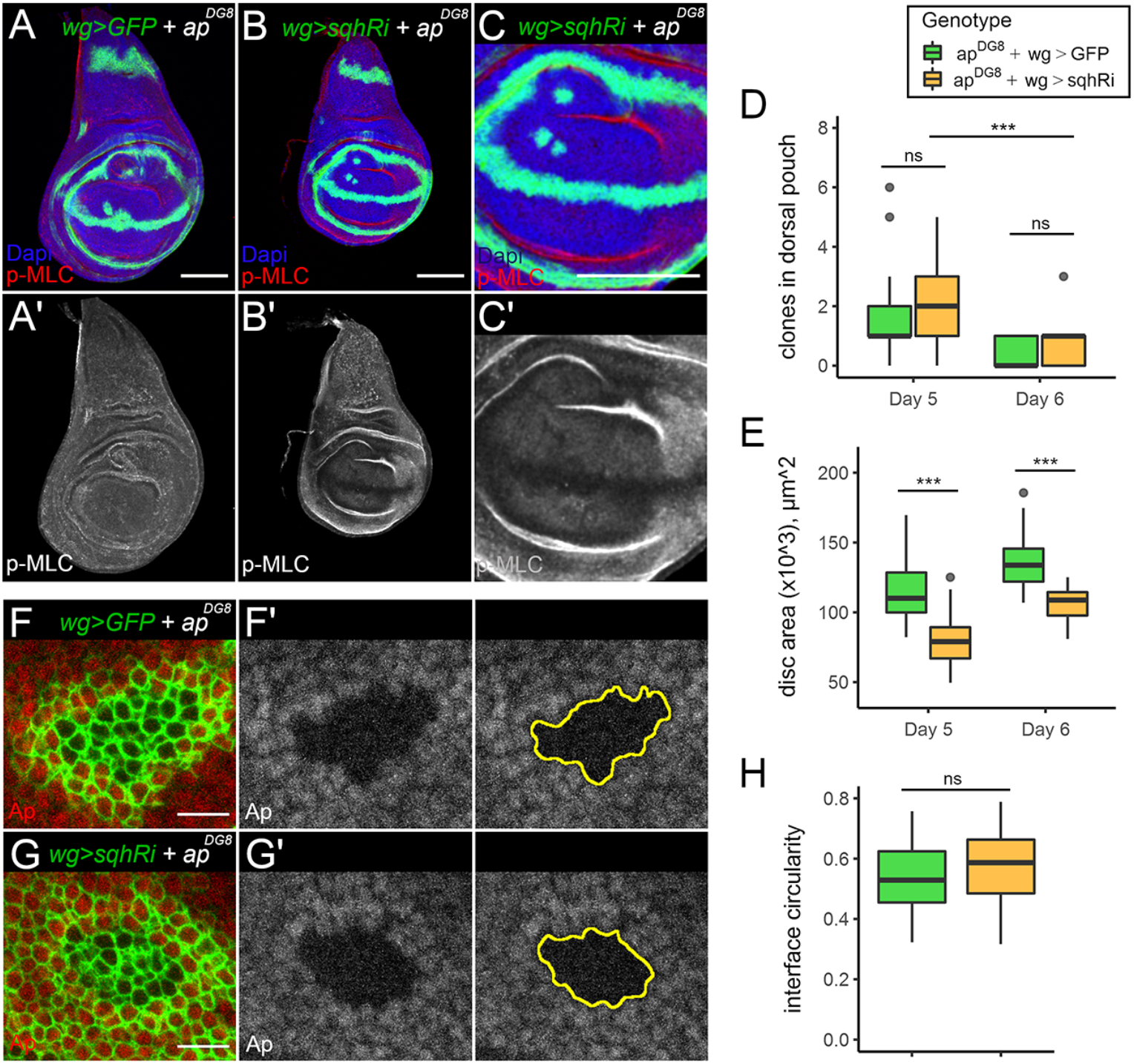
Knocking down Myosin light chain around *ap^DG8^* clones does not affect clone recovery and circularity. **A-C’)** day 5 wing discs bearing unmarked clones that are mutant for *ap^DG8^* and expressing GFP (A) or GFP and *sqh-RNAi* (B, C’) under *wg::Gal4* control. Remaining clones in the pouch are detectable with the ectopic *wg::Gal4* driven *GFP* induced around them. Panel C shows the pouch region of the disc in panel B. Nuclei are marked with DAPI (blue) and p-MLC staining is shown in red (top panels) or in grey (lower panels). Scale bars in (A-C) are 100μm. Dorsal is up, anterior is to the left in all panels. **D-E)** Quantification of the number of remaining clones in the dorsal pouch (D) and disc areas (E) in indicated genotypes. At least 36 discs per genotype for day 5 and at least 15 discs per genotype for day 6 were considered for the analysis. Expression of *sqh-RNAi* causes growth delay, and further elimination happens between the two time points. **F-G’)** Representative *ap^DG8^* clones expressing GFP (F) or GFP and *sqh-RNAi* (G) under *wg::Gal4* control. F’ and G’ show Ap antibody staining in grey and estimated clone boundaries based on this staining in yellow. **H)** Quantification of interface circularity of the remaining *ap^DG8^* clones with (yellow) or without (green) *wg::Gal4* driven *sqh-RNAi* in the dorsal pouch. 40 clones were measured for each genotype. Scale bars in (F-G) are 10μm.

## DISCUSSION

The forces produced by actomyosin cables are essential and were implicated in many biological processes, like tissue closure, wound healing, tissue extension, tube formation, compartment organization and cell elimination [4, 10, 47]. The importance of actomyosin cables in some of these processes has been recently revised [48, 49]. In this study, we challenge the model, which attributes a central role for interface contractility mediated by actomyosin filaments in identification and elimination of aberrantly specified cells.

The interactions of differently fated cells form boundaries that restrict cell mixing and maintain tissue organization. Previous work has shown that accumulation of actomyosin filaments and increased cell bound tension are vital to boundary function [1–3, 29]. While this physiological mechanism works well in separating two similarly sized cell populations (such as two compartments), it could be problematic when the populations differ in size, and one encloses the other. The cells of the underrepresented group encircled by a highly tense boundary experience compression [4]. Mechanical cell deformation can in turn induce cell death and elimination, possibly, via downregulation of the EGFR/ERK signaling pathway [50]. This model elegantly explains how misspecified cells are detected and why apoptosis is associated with misspecified cells, but not with compartment boundaries.

In our study we used cells with aberrant dorsoventral identity to test this model. We demonstrated that the interface between Ap positive and Ap negative cell groups is indeed enriched for both F-actin and Myosin II cables (Fig. 3). The interface is highly smooth and round (Fig. 2A-D), suggesting increased tension. Moreover, the cells of underrepresented populations show obvious apical constrictions (Fig. 2E). All these observations are in favor of the model and are consistent with previous studies [1, 4, 25]. However, when we tested the involvement of non-muscle Myosin II in the process, we found that it was largely dispensable.

We have assessed the importance of Myosin-driven tension in the elimination of misspecified cell populations using three different approaches: tissue-wide reduction of Myosin II function (Fig. 7B), inhibition in misspecified cells only (Fig. 7C) or specifically in cells flanking the interface on either side (Fig. 7D). None of the approaches showed a strong rescue effect on misspecified cell clones; moreover, Myosin depletion did not hinder their tendency to round up and form a smooth interface with wt neighbors. These results have several possible explanations. Technical limitations of the approaches used are one possibility. As non-muscle Myosin II is an essential molecule, and its function is required for cytokinesis, it is impossible to remove it completely from cells that are expected to divide and form clones. In our study, a hypomorphic mutant combination of Myosin heavy chain (*zip^Ebr^/zip^2^*) or an RNAi against the Myosin light chain (s*qhRi*) were used to reduce Myosin II levels. Notably, the depletion of Myosin with *zip^Ebr^/zip^2^* was less efficient than that of *sqhRi* and some junctional Myosin was easily detected (Fig. 7B-B’). Therefore, it is possible that the residual Myosin in the *zip^Ebr^/zip^2^* cells could generate enough elevated tension along the interface to cause clone rounding and elimination. Myosin depletion in misspecified cells using *sqhRi* does alter clone circularity and elimination efficiency, but this effect was extremely minor (Fig. 5F, 5K). The previous work demonstrated that clone separation depends on the relative difference in junctional tension between clone bulk and its boundary, meaning that the reduction of tension within the clonal cells is enough to cause clone separation [51]. However, our data shows that reduction of tension by *sqhRi* in cell clones does not lead to clone rounding (Fig. 5I, 5K). Moreover, *sqhRi* clones dissociate and their cells mix with wt cells more readily (Fig. S1A, S1C). This is likely because the removal of Myosin from one side of the junction (in clonal cell) cannot be compensated for by Myosin accumulation from the opposite side of the junction (wt cell), thus the resulting tension of the interfacial junction is also relatively low. However, in the context of misspecified cells, actomyosin levels at interfacial junctions are strongly elevated compared to those in either clone bulk or surrounding wt cells (Fig. 3A-B’ and 7A). Hence, if actomyosin accumulation along the shared junction of two contacting cells occurs independently, the reduction of Myosin only in clonal cells could be inadequate in abolishing the increased interfacial tension generated from the wt side. This could explain why the rescue effect in the second approach is too mild. This potential problem was overcome in the third approach, where Myosin function was depleted in flanking cells from either side of the boundary (Fig. 7D). Although the strategy by which we measured and quantified the efficiency of clone elimination and clone circularity was different from the previous two, the results clearly demonstrate that rounding and elimination of misspecified clones do not rely on Myosin accumulation along the boundary between differently specified cells. Altogether, our results suggest that in the context of aberrantly specified cell clone elimination, the interfacial contractility is either not as important as it had been previously presumed, or the interfacial tension is Myosin independent.

**Figure 7:**
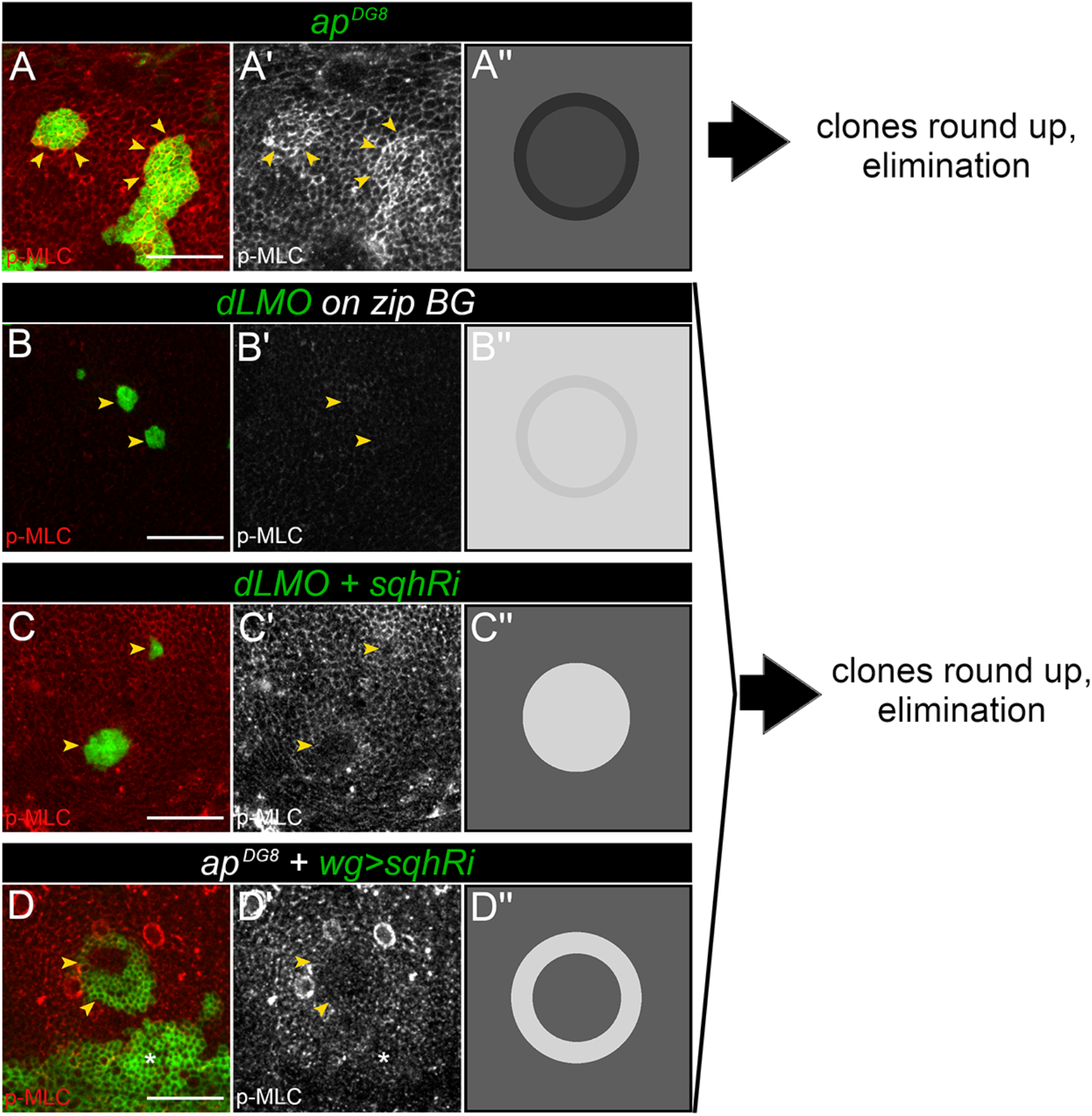
Interfering with Myosin function with different approaches is not sufficient to abolish elimination of the misspecified cell clones. **A-D)** Representative images and schematics (A’’-D’’) to summarize and compile the data presented in Figure 3 (A), Figure 4 (B), Figure 5 (C), and Figure 6 (D). **A)** GFP-marked *ap^DG8^* clones **B)** Clones expressing *dLMO* and GFP (green) in *zip^Ebr/2^* mutant background **C)** Clones expressing *dLMO* and *sqh-RNAi* (green), **D)** *ap^DG8^* clone with *wg::Gal4* driven *sqh-RNAi* (green), Wg induction around the clone was used to downregulate Myosin function at the clone boundary. The GFP positive area below the clone (white star) corresponds to the DV boundary. Panels (A -D) show clone of corresponding genotype in green and p-MLC staining in red. Panels (A’-D’) show p-MLC staining alone in grey. Yellow arrowheads point to the clone borders. Panels (A’’-D’’) schematize Myosin II distribution for each approach. Scale bars represent 20μm in all panels.

In conclusion, we revised the model that attributes the main role of identification and elimination of aberrantly specified cells to interfacial supra-cellular contractility driven by actomyosin cables. Partial rescue effect of *sqhRi* co-expression in misspecified cell clones suggests that Myosin II mediated tension contributes to cell separation and elimination, but its contribution is rather minor. Thus, we have ruled out a decisive role for Myosin II driven tension in elimination of mispositioned cell clusters.

## MATERIALS AND METHODS

### *Drosophila* strains

The following *Drosophila* stocks were used: *ap^DG8^*, *FRT^f00878^* [52] and *UAS-sqh-GFP* [53] (kindly provided by Markus Affolter), *UAS-Ap* and *UAS-dLMO* [54] (kindly provided by Marco Milan), *zip^Ebr^* [55] and *zip^2^* [56] (kindly provided by Romain Levayer), *UAS-sqhRNAi* (GD7917) (obtained from Vienna Drosophila Resource Center (VDRC)), *wg::Gal4* [46] (kindly provided by Jean-Paul Vincent). All crosses were kept on standard media at 25°C. Flippase expression was induced by a heat shock at 37°C. The detailed fly genotypes and experimental conditions are presented in Table S1.

### Immunohistochemistry

Imaginal discs were prepared and stained using standard procedures. Briefly, larvae were dissected in cold PBS and fixed in 4% paraformaldehyde (PFA) in PBS for 20 min. Washes were performed in PBS + 0.03% Triton X-100 (PBT) and blocking in PBT + 2% normal donkey serum (PBTN). Samples were incubated with primary antibodies overnight at 4°C. The primary antibodies used: mouse anti-Wingless (1:2000, was deposited to the DSHB by Cohen, S.M. (DSHB Hybridoma Product 4D4)), rat anti-DE-Cadherin (1:30, was deposited to the DSHB by Uemura, T. (DSHB Hybridoma Product DCAD2)), rabbit anti-phospho-Myosin light chain 2 (Ser19) (1:50, Cell Signaling Technology #3671), rabbit anti-Ap (1:1000, described in [52]). Incubation with secondary antibodies were at room temperature, for 2hr. The secondary antibodies used: anti-mouse Alexa 568 (1:700, ThermoFisher) and Alexa 633 (1:700, ThermoFisher), anti-rat Cy3 (1:300, Jackson ImmunoResearch) anti-rabbit Alexa 568 (1:600, ThermoFisher) and Alexa 633 (1:600, ThermoFisher). Discs were mounted in Vectashield antifade mounting medium with Dapi (Vector Laboratories). For F-actin staining Phalloidin-Tetramethylrhodamine B (Fluka #77418) was added during incubation with secondary antibodies at the concentration 0.3 μM.

### TUNEL assay

For the TUNEL assay, *In Situ* Cell Death Detection kit, TMR red (Roche) was used. Larvae were dissected in cold PBS and fixed in 4% PFA for 1hr at 4°C. Samples were washed in PBT and blocked in PBTN for 1 hr. Next, samples were incubated with primary antibodies overnight at 4°C and with secondary antibodies for 4hr at 4*°*C. After washing the tissues were incubated in PBTN overnight at 4°C. Then, samples were permeabilized in 100 mM sodium citrate supplemented with 0.1% Triton X-100 and incubated in 50 μl of TUNEL reaction mix (prepared according to the recipe from the kit) for 2hr at 37°C in the dark. After this step, the samples were washed in PBT for 30 min and mounted in Vectashield antifade mounting medium with Dapi (Vector Laboratories).

### Image acquisition

Image stacks of wing discs were acquired on a Zeiss LSM880 confocal microscope using 20x, 40x, and 63x objectives. The pinhole was 1 airy unit. The intervals between Z-sections were 1μm thick (for 20x) and 0.4 - 0.5μm thick (for 40x and 63x).

### Image analysis

Clone area measurements were done using ImageJ image processing platform. Image stacks were projected using maximum projections. All z-slices were included in the projection except the ones containing signal from the peripodial membrane. Dorsal compartment was selected manually based on Wingless staining. For clone detection, Gaussian Blur filter (sigma = 2.0) was applied and clones were detected using intensity based thresholding.

Clone circularity and cell area measurements were done on images taken with 63x objective. Individual clones were selected and processed separately. For experiments presented in Fig. 2, 4 and 5, clone perimeter and cell area were measured using image analysis toolkit Epitools [45]. Two to eight z-slices from the apical part of the stack were projected using selective plane projection (built-in algorithm in Epitools). Cell outlines were detected based on DE-Cadherin staining. Cell segmentation was done using MATLAB-based analysis framework of Epitools. Clones were detected using the GFP signal. Clone area, perimeter and number of clonal cells, as well as cell areas within the clone and in surrounding bulk were obtained using CELL_CLONE and CELL_AREA Epitools modules (cellGraph v0.9.1.0) respectively. Circularity was quantified using formula 4π x area/perimeter^2^. The heatmaps of cell areas scale from 0.2μm^2^ (deep red) to 12μm^2^ (light green) across all experiments. For experiments presented in Fig. 6, the clones were detected and the circularity was measured using ImageJ. Ten z-slices from the center of the stack were projected using maximum projection. Gaussian Blur filter (sigma 3.5) was applied and the mutant clones were identified based on loss of Apterous staining using intensity thresholding (low intensity).

Mixing index and contact length (Fig. S1) were quantified using the dataset presented in Fig. 5. The number of clone border cells, number of wt contacting cells and clone perimeters were obtained from CELL_CLONE Epitools module.

### Statistical analysis

Data processing and statistical analysis were done in R, v3.5.0. Conditions were compared using two-sample t-test. Comparisons with a p-value ≥ 0.05 were marked as “ns” (non-significant); p-value < 0.05 - “*”; p-value < 0.01 –“**”; p-value < 0.001 –“***”. Note, for circularity measurements in Fig. 5K, as well as mixing index and contact length measurements in Fig. S1C and S1D, 12 out of 47 sqhRNAi data points were removed from the analysis. Those corresponded to small GFP positive groups of cells (with 1-10 cells size), located close to bigger clones and thus, they were presumed to be split from the bigger clones.

## Acknowledgements

Confocal microscopy was performed at the Cardiff University Bioimaging Research Hub, we thank the team for their support. We are grateful to the amazing *Drosophila* community for providing reagents and for the countless discussions. Special thanks to Davide Heller for creating the CELL_CLONE module in EpiTools upon our request. We thank Vincent Dion and members of the Hamaratoglu Lab for comments on the manuscript.

---

**Figure S1.**
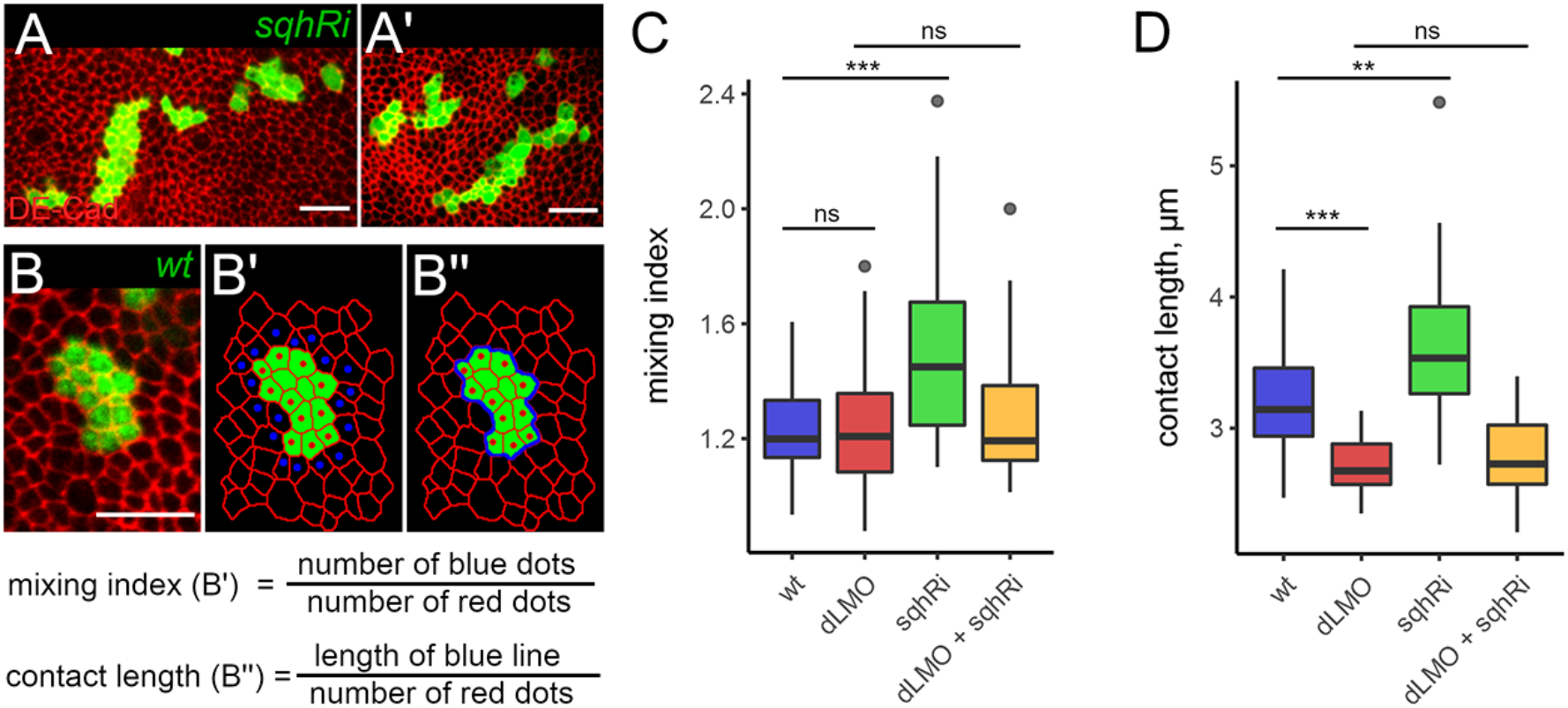
(related to Figure 5): Cell clones expressing *sqh-RNAi* disperse and mix with their wild-type neighbours. **A-A’)** Examples of cell clones expressing *sqh-RNAi* (green) stained for DE-Cad (red). Such clones display a more dispersed morphology than wild-type clones. **B)** A GFP-marked (green) wild-type clone in a disc stained for DE-Cad (red). **B’-B’’)** Schematics explain the logic used for driving the mixing index and contact length formulas shown below. Red dots show clonal cells touching the boundary, whereas blue dots mark wild-type cells touching the boundary. Clone outline (perimeter of the clone) is shown as a blue line in (B’’). **C-D)** Quantification of the mixing index (C) and contact length (D) for clones of the indicated genotypes. Minimum 30 clones per genotype were analysed. Cells in *sqh-RNAi* clones contact a higher number of surrounding cells and these contacts are longer. *dLMO* expression on its own or along with *sqh-RNAi* reduces the contact length with the surrounding cells. Scale bars represent 10μm in all panels.

**Figure S2.**
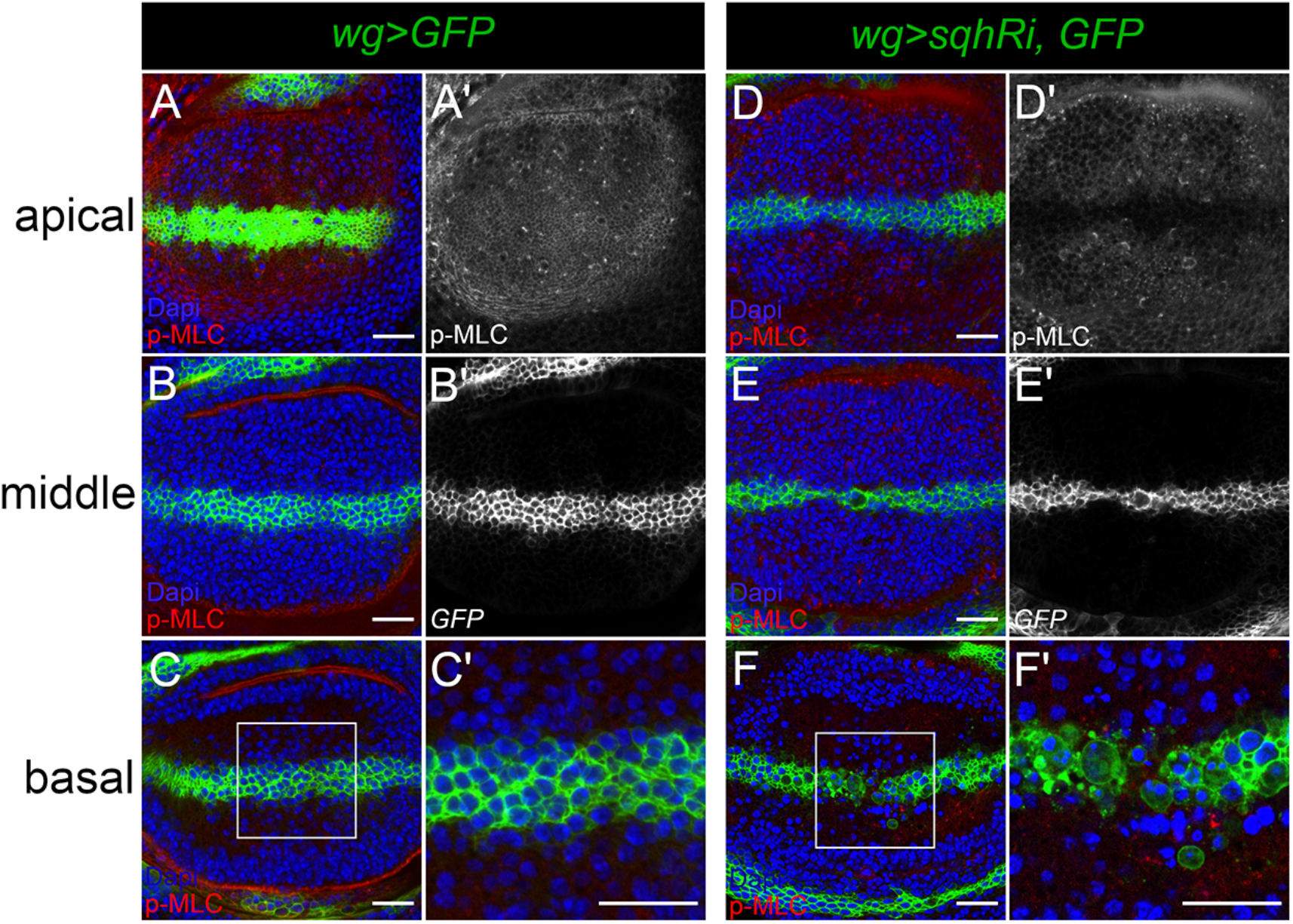
(related to Figure 6): *sqh-RNAi* expression at the DV boundary undermines integrity of the boundary. **A-B)** Pouch regions from representative discs expressing GFP (green) (left) or GFP and *sqh-RNAi* (right) under *wg::Gal4* control, stained for DAPI (blue) and p-MLC (red). Optical sections at three different levels are shown: apical, middle and basal. C’ and F’ shows zoomed versions of white boxes in C and F. A gap in the boundary and dying cells are visible at the basal side upon *sqh* depletion. Scale bars represent 20μm in all panels. Dorsal is up, anterior is to the left in all panels.

**Table S1:**
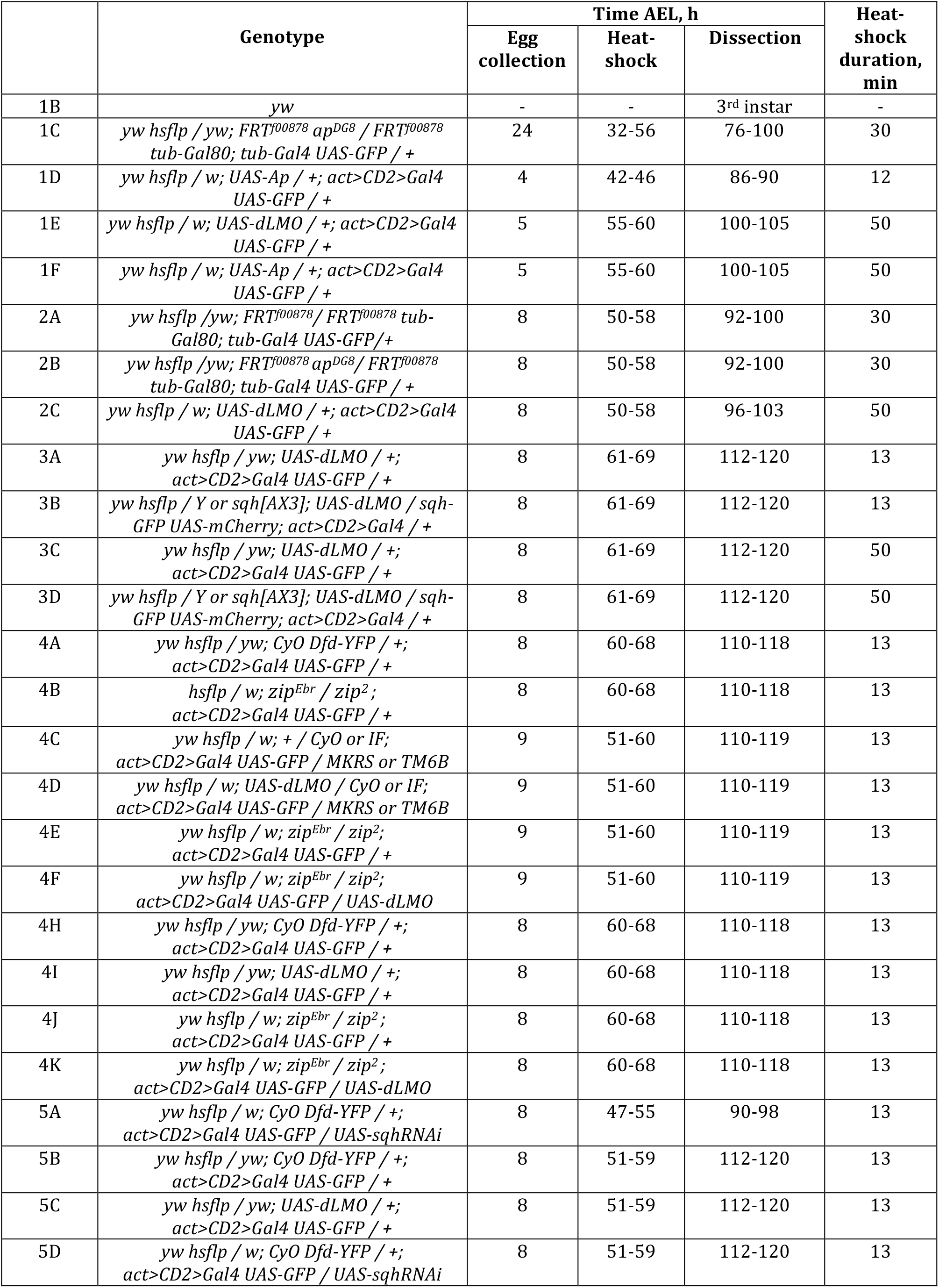
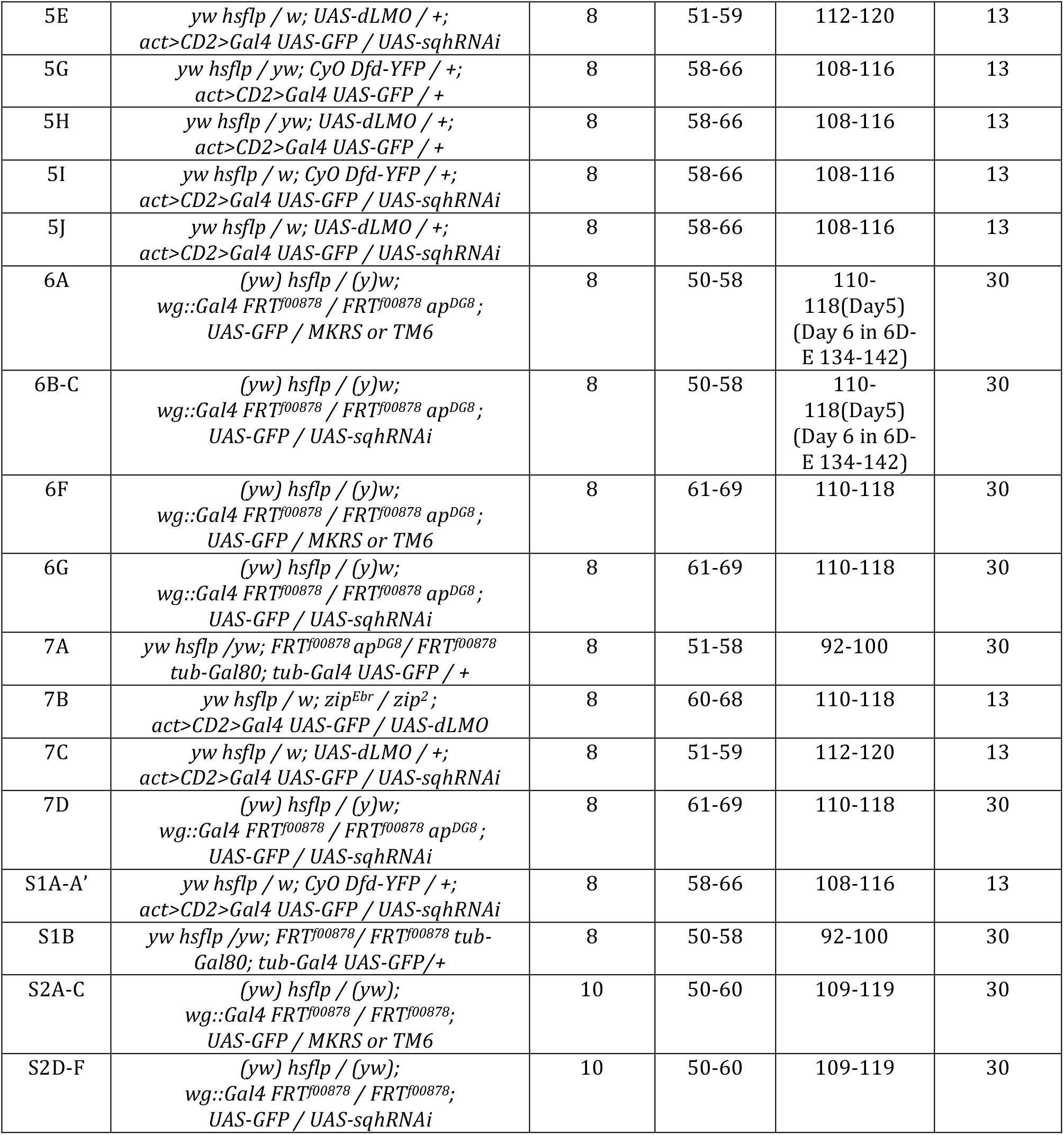
Genotypes and experimental set-up for each figure panel.

## Notes

### Competing Interest Statement

The authors have declared no competing interest.

### Summary of Updates

TableS1, which shows detailed fly genotypes and experimental set-up, is included in the revised version.

## REFERENCES

1. Major, R.J. and K.D. Irvine, Localization and requirement for Myosin II at the dorsal-ventral compartment boundary of the Drosophila wing. Dev Dyn, 2006. 235(11): p. 3051–8.

2. Landsberg, K.P., et al., Increased cell bond tension governs cell sorting at the Drosophila anteroposterior compartment boundary. Curr Biol, 2009. 19(22): p. 1950–5.

3. Aliee, M., et al., Physical mechanisms shaping the Drosophila dorsoventral compartment boundary. Curr Biol, 2012. 22(11): p. 967–76.

4. Bielmeier, C., et al., Interface Contractility between Differently Fated Cells Drives Cell Elimination and Cyst Formation. Curr Biol, 2016. 26(5): p. 563–74.

5. Adachi-Yamada, T., et al., Distortion of proximodistal information causes JNK-dependent apoptosis in Drosophila wing. Nature, 1999. 400(6740): p. 166–9.

6. Adachi-Yamada, T. and M.B. O’Connor, Morphogenetic apoptosis: a mechanism for correcting discontinuities in morphogen gradients. Dev Biol, 2002. 251(1): p. 74–90.

7. Klipa, O. and F. Hamaratoglu, Cell elimination strategies upon identity switch via modulation of apterous in Drosophila wing disc. PLoS Genet, 2019. 15(12): p. e1008573.

8. Beira, J.V. and R. Paro, The legacy of Drosophila imaginal discs. Chromosoma, 2016. 125(4): p. 573–92.

9. Ruiz-Losada, M., et al., Specification and Patterning of. J Dev Biol, 2018. 6(3).

10. Wang, J. and C. Dahmann, Establishing compartment boundaries in Drosophila wing imaginal discs: An interplay between selector genes, signaling pathways and cell mechanics. Semin Cell Dev Biol, 2020. 107: p. 161–169.

11. Steinberg, M.S., Reconstruction of tissues by dissociated cells. Some morphogenetic tissue movements and the sorting out of embryonic cells may have a common explanation. Science, 1963. 141(3579): p. 401–8.

12. Foty, R.A. and M.S. Steinberg, The differential adhesion hypothesis: a direct evaluation. Dev Biol, 2005. 278(1): p. 255–63.

13. Tepass, U., D. Godt, and R. Winklbauer, Cell sorting in animal development: signalling and adhesive mechanisms in the formation of tissue boundaries. Curr Opin Genet Dev, 2002. 12(5): p. 572–82.

14. Brodland, G.W., The Differential Interfacial Tension Hypothesis (DITH): a comprehensive theory for the self-rearrangement of embryonic cells and tissues. J Biomech Eng, 2002. 124(2): p. 188–97.

15. Lecuit, T. and P.F. Lenne, Cell surface mechanics and the control of cell shape, tissue patterns and morphogenesis. Nat Rev Mol Cell Biol, 2007. 8(8): p. 633–44.

16. Fagotto, F., The cellular basis of tissue separation. Development, 2014. 141(17): p. 3303–18.

17. Milán, M., et al., The LRR proteins capricious and Tartan mediate cell interactions during DV boundary formation in the Drosophila wing. Cell, 2001. 106(6): p. 785–94.

18. Milán, M. and S.M. Cohen, A re-evaluation of the contributions of Apterous and Notch to the dorsoventral lineage restriction boundary in the Drosophila wing. Development, 2003. 130(3): p. 553–62.

19. Rauskolb, C., T. Correia, and K.D. Irvine, Fringe-dependent separation of dorsal and ventral cells in the Drosophila wing. Nature, 1999. 401(6752): p. 476–80.

20. O’Keefe, D.D. and J.B. Thomas, Drosophila wing development in the absence of dorsal identity. Development, 2001. 128(5): p. 703–10.

21. Becam, I., et al., Notch-mediated repression of bantam miRNA contributes to boundary formation in the Drosophila wing. Development, 2011. 138(17): p. 3781–9.

22. Milán, M., L. Pérez, and S.M. Cohen, Boundary formation in the Drosophila wing: functional dissection of Capricious and Tartan. Dev Dyn, 2005. 233(3): p. 804–10.

23. Micchelli, C.A. and S.S. Blair, Dorsoventral lineage restriction in wing imaginal discs requires Notch. Nature, 1999. 401(6752): p. 473–6.

24. Milán, M. and S.M. Cohen, Temporal regulation of apterous activity during development of the Drosophila wing. Development, 2000. 127(14): p. 3069–78.

25. Major, R.J. and K.D. Irvine, Influence of Notch on dorsoventral compartmentalization and actin organization in the Drosophila wing. Development, 2005. 132(17): p. 3823–33.

26. Michel, M., et al., The Selector Gene apterous and Notch Are Required to Locally Increase Mechanical Cell Bond Tension at the Drosophila Dorsoventral Compartment Boundary. PLoS One, 2016. 11(8): p. e0161668.

27. Becam, I. and M. Milán, A permissive role of Notch in maintaining the DV affinity boundary of the Drosophila wing. Dev Biol, 2008. 322(1): p. 190–8.

28. Umetsu, D., et al., Local increases in mechanical tension shape compartment boundaries by biasing cell intercalations. Curr Biol, 2014. 24(15): p. 1798–805.

29. Monier, B., et al., An actomyosin-based barrier inhibits cell mixing at compartmental boundaries in Drosophila embryos. Nat Cell Biol, 2010. 12(1): p. 60–9.

30. Rudolf, K., et al., A local difference in Hedgehog signal transduction increases mechanical cell bond tension and biases cell intercalations along the Drosophila anteroposterior compartment boundary. Development, 2015. 142(22): p. 3845–58.

31. Morata, G. and P.A. Lawrence, Control of compartment development by the engrailed gene in Drosophila. Nature, 1975. 255(5510): p. 614–7.

32. Diaz-Benjumea, F.J. and S.M. Cohen, Interaction between dorsal and ventral cells in the imaginal disc directs wing development in Drosophila. Cell, 1993. 75(4): p. 741–52.

33. Blair, S.S., et al., The role of apterous in the control of dorsoventral compartmentalization and PS integrin gene expression in the developing wing of Drosophila. Development, 1994. 120(7): p. 1805–15.

34. Milán, M., L. Pérez, and S.M. Cohen, Short-range cell interactions and cell survival in the Drosophila wing. Dev Cell, 2002. 2(6): p. 797–805.

35. Dahmann, C. and K. Basler, Opposing transcriptional outputs of Hedgehog signaling and engrailed control compartmental cell sorting at the Drosophila A/P boundary. Cell, 2000. 100(4): p. 411–22.

36. Baena-Lopez, L.A. and A. García-Bellido, Control of growth and positional information by the graded vestigial expression pattern in the wing of Drosophila melanogaster. Proc Natl Acad Sci U S A, 2006. 103(37): p. 13734–9.

37. Villa-Cuesta, E., E. González-Pérez, and J. Modolell, Apposition of iroquois expressing and non-expressing cells leads to cell sorting and fold formation in the Drosophila imaginal wing disc. BMC Dev Biol, 2007. 7: p. 106.

38. Shen, J., C. Dahmann, and G.O. Pflugfelder, Spatial discontinuity of optomotor-blind expression in the Drosophila wing imaginal disc disrupts epithelial architecture and promotes cell sorting. BMC Dev Biol, 2010. 10: p. 23.

39. Moreno, E., K. Basler, and G. Morata, Cells compete for decapentaplegic survival factor to prevent apoptosis in Drosophila wing development. Nature, 2002. 416(6882): p. 755–9.

40. Giraldez, A.J. and S.M. Cohen, Wingless and Notch signaling provide cell survival cues and control cell proliferation during wing development. Development, 2003. 130(26): p. 6533–43.

41. Johnston, L.A. and A.L. Sanders, Wingless promotes cell survival but constrains growth during Drosophila wing development. Nat Cell Biol, 2003. 5(9): p. 827–33.

42. Gibson, M.C. and N. Perrimon, Extrusion and death of DPP/BMP-compromised epithelial cells in the developing Drosophila wing. Science, 2005. 307(5716): p. 1785–9.

43. Shen, J. and C. Dahmann, Extrusion of cells with inappropriate Dpp signaling from Drosophila wing disc epithelia. Science, 2005. 307(5716): p. 1789–90.

44. Widmann, T.J. and C. Dahmann, Wingless signaling and the control of cell shape in Drosophila wing imaginal discs. Dev Biol, 2009. 334(1): p. 161–73.

45. Heller, D., et al., EpiTools: An Open-Source Image Analysis Toolkit for Quantifying Epithelial Growth Dynamics. Dev Cell, 2016. 36(1): p. 103–116.

46. Alexandre, C., A. Baena-Lopez, and J.P. Vincent, Patterning and growth control by membrane-tethered Wingless. Nature, 2014. 505(7482): p. 180–5.

47. Röper, K., Supracellular actomyosin assemblies during development. Bioarchitecture, 2013. 3(2): p. 45–9.

48. Pasakarnis, L., et al., Amnioserosa cell constriction but not epidermal actin cable tension autonomously drives dorsal closure. Nat Cell Biol, 2016. 18(11): p. 1161–1172.

49. Ochoa-Espinosa, A., et al., Myosin II is not required for. Development, 2017. 144(16): p. 2961–2968.

50. Moreno, E., et al., Competition for Space Induces Cell Elimination through Compaction-Driven ERK Downregulation. Curr Biol, 2019. 29(1): p. 23–34.e8.

51. Bosveld, F., et al., Modulation of junction tension by tumor suppressors and proto-oncogenes regulates cell-cell contacts. Development, 2016. 143(4): p. 623–34.

52. Bieli, D., et al., Establishment of a Developmental Compartment Requires Interactions between Three Synergistic Cis-regulatory Modules. PLoS Genet, 2015. 11(10): p. e1005376.

53. Royou, A., et al., Reassessing the role and dynamics of nonmuscle myosin II during furrow formation in early Drosophila embryos. Mol Biol Cell, 2004. 15(2): p. 838–50.

54. Milán, M. and S.M. Cohen, Regulation of LIM homeodomain activity in vivo: a tetramer of dLDB and apterous confers activity and capacity for regulation by dLMO. Mol Cell, 1999. 4(2): p. 267–73.

55. Halsell, S.R., B.I. Chu, and D.P. Kiehart, Genetic analysis demonstrates a direct link between rho signaling and nonmuscle myosin function during drosophila morphogenesis. Genetics, 2000. 156(1): p. 469.

56. Young, P.E., et al., Morphogenesis in Drosophila requires nonmuscle myosin heavy chain function. Genes Dev, 1993. 7(1): p. 29–41.

